# Prediction and validation of host cleavage targets of SARS-CoV-2 3C-like protease

**DOI:** 10.1101/2022.01.17.476677

**Authors:** Nora Yucel, Silvia Marchiano, Evan Tchelepi, Germana Paterlini, Quentin McAfee, Nehaar Nimmagadda, Andy Ren, Sam Shi, Charles Murry, Zoltan Arany

**Author notes:** Corresponding: Zoltan Arany MD PhD Professor in Medicine Perelman School of Medicine University of Pennsylvania, Smilow Center for Translational Research 11th floor, 3400 Civic Blvd, Philadelphia 19104. Administrative assistant: Emily Romick.

## Abstract

How SARS-CoV-2 causes the observed range of clinical manifestations and disease severity remains poorly understood. SARS-CoV-2 encodes for two proteases (3CLPro and PLPro), vital for viral production, but also promiscuous with respect to host protein targets, likely contributing to the range of disease. Pharmacological inhibition of the 3C-like3 protease has revealed remarkable reduction in hospitalization and death in phase 2/3 clinical studies. However, the mechanisms responsible for the pathology mediated by those proteases are still unclear. In this study, we develop a bioinformatic algorithm, leveraging experimental data from SARS-CoV, to predict host cleavage targets of the SARS-CoV-2 3C-like protease, or 3CLPro. We capture targets of the 3CL protease described previously for SARS-CoV, and we identify hundreds of new putative targets. We experimentally validate a number of these predicted targets, including the giant sarcomeric protein Obscurin, and show that expression of 3CL protease alone recapitulates the sarcomeric disorganization seen by SARS-CoV-2 infection of hiPSC-derived cardiomyocytes. Our data provide a resource to identify putative host cleavage targets of 3CL protease that contribute to mechanisms and heterogeneity of disease in COVID-19 and future coronavirus outbreaks.

## Introduction

COVID-19 continues to be a leading cause of death and morbidity across the world since the initial outbreak in Wuhan, China December 2019^1^. How SARS-CoV-2, the causative agents of COVID19, leads to its wide range of disease manifestations remains incompletely understood. In addition to lung damage, SARS-CoV-2 infection can also cause kidney damage, clotting disorders, loss of taste and smell, cognitive dysfunction, muscle atrophy, and cardiac dysfunction ^2–5^. In addition, long-lasting COVID-19 symptoms have been reported in patients up to a year after initial illness, including fatigue, shortness of breath, brain fog, and elevated heart rate ^6, 7^. It remains unclear how SARS-CoV-2 affects differently the multiple organs and cell types involved. Nor it is known what host characteristics determine who will develop severe COVID-19, which organs will be affected, or whether long COVID will ensue. A deeper mechanistic understanding of virus-host interactions is thus needed.

Various studies have identified host interaction partners of many SARS-CoV-2 proteins, including the spike, envelope, nucleocapsid, and membrane proteins^8–10^. These interactions have various consequences, including suppression of innate immune response, suppression of apoptosis, and reprogramming of host transcription and translation. In addition to protein-protein interactions, important virus-host interactions can be caused by enzymatic cleavage of host proteins by viral proteases. For example, myocardial dysfunction following infection by coxsackie CVB3 virus can in part be ascribed to cleavage of dystrophin protein by the viral protease 2A^11, 12^; enteroviral 3C proteases can cleave host NLRP1 to trigger inflammasome activation ^13^; HIV-1 protease mediates apoptosis by cleaving host procaspase 9 and Bcl2^14, 15^; and the Zika virus nsP2 cysteine protease can cleave host proteins SFRP1 NT5M, and FOXG1^16^.

SARS-CoV-2 encodes for two proteases, a papain-like protease (PLPro) and the 3C-like protease (3CLPro, also knowns as main Protease, MPro, or NSP5) . These proteases are highly conserved across coronavirus species and are absolutely required for viral replication^17–20^. Both proteases are thus actively being investigated as targets for antivirals ^21^. Recent Phase2/3 clinical trial results with PF-07323332, a 3CL inhibitor, administered in combination with ritonavir, revealed 89% reduction in hospitalization with COVID-19 ^22^. Both PLPro and 3CLPro are generated via autocatalytic cleavage from the overlapping ORF1a and ORF1ab polyproteins, the first translation products following SARS-CoV-2 infection. The ORF1a/ab polyproteins encode 16 non-structural proteins (NSP1-16) that build the viral replication machinery (**Fig 1A**). The nonstructural proteins are liberated from the ORF1a/ab polyprotein through cleavage by PLPro and 3CLPro, encoded by NSP3 and NSP5, respectively. Following this processing PLPro remains bound to the endoplasmic reticulum membrane, while 3CLPro cleaves itself free, giving it access to the full cytosolic compartment. The impact of these proteases on the host proteome, and in particular the 3CLPro, remains poorly defined. Targeted screening of 300 interferon-stimulated proteins in cell lines overexpressing SARS-CoV-2 3CLPro identified RNF20 as a target of 3CLpro^23^. In a different screening of 71 immune pathway-related proteins, IRF3 was identified as target of PLpro and NLRP12 and TAB1 as targets of 3CLpro, suggesting the role of those proteases in the innate immune response to the virus ^24^.

**Fig 1.**
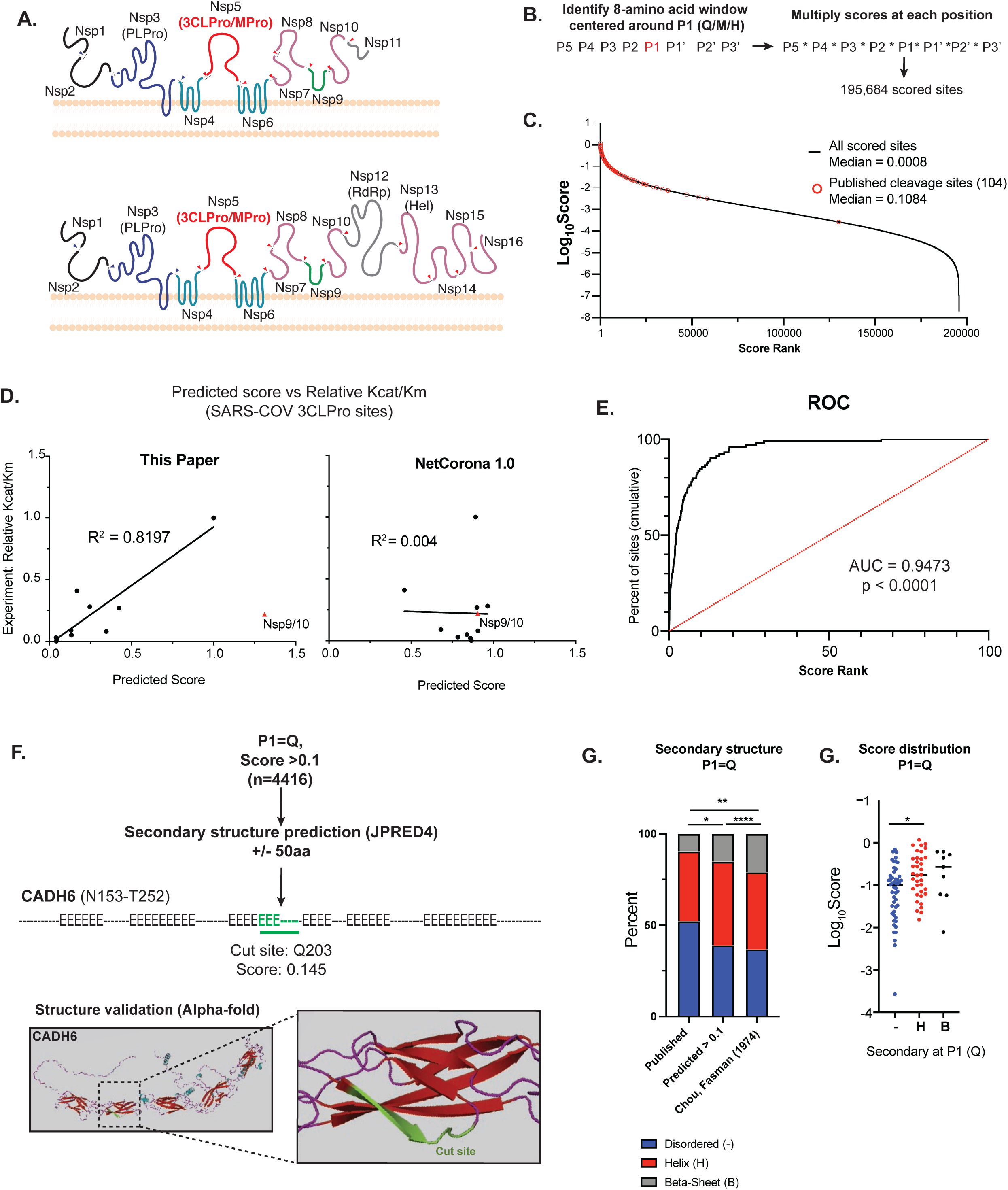
Bioinformatic prediction of SARS-COV2 3CL human protein targets. A) Diagram of endogenous function of the SARS-CoV-2 3C-like protease (3CLPro). 3CLPro cleaves at 11 sites within the 2 large polypeptides pp1a and pp1ab generated from the overlapping reading frames ORF1a and ORF1ab (respectively). Cleavage by 3CLPro, as well as the other viral protease papain-like protease (PLPro) liberates the nonstructural proteins (NSPs) that are required for viral transcription, replication, suppression of host immune responses and suppression of host gene expression. B) Work-flow for bioinformatic identification and scoring of putative 3CLPro cleavage sites withing the human proteome. Scores for each position along the cleavage site (P5-P3’) were obtained from experimental data from SARS-CoV 3CLPro (**Chuck, et al 2010**). First P1 positions were identified (Q, M or H), and overall cleavage score generated by multiplying scores for each amino acid around P1. Within each 8-amino acid window, any position that contained an amino acid that scored a “ND” (no cleavage detected) in **Chuck, et al** resulted in a score of 0. Overall, 195,684 scored cleavage sites (>0) were detected across the human proteome. C) Distribution of all scores (Log_10_Score). Scores of published cleavage sites detected by our prediction are highlighted in red. D) Correlation of predicted score with experimentally-derived K_cat_/K_m_ values for SARS-COV 3CLPro **(Grums-Tokars, 2008).** Scores generated in this study are shown on the left, and scores generated by NetCorona1.0 (**Kiemer, 2004**). Shown on the right. For R^2^ calculations, the cleavage site between NSP9 and NSP10 for both graphs. E) Receiver operator curve analysis to assess predictive power of bioinformatic scoring based on scores of published cleavage sites. Cumulative percentage of scores captured plotted vs score rank (highest score = 1 to lowest score = 100), and area under the curve (AUC) captured. 95% confidence interval determined by Wilson/Brown method. F) Secondary structure analysis of high scoring sites (>0.1) with P1 = Q. To increase secondary structure accuracy, a 100aa window centered around P1 (Q) was identified. Resulting 100aa peptides were analyses by JPRED4 to predict secondary structure ( “-“ = unstructured, “H” = alpha-helix, “E” = beta-sheet). When available, candidate cleavage sites were verified by alpha-Fold structure. Highlighted is a predicted site in Cadherin-6 (CADH6). G) Fraction of each P1 (Q) that lies in each type of secondary structure (unstructured, alpha-helix, beta sheet). Comparisons shown for predicted cleavage sites with score > 0.1 vs published cleavage sites (all scores) vs published secondary structure distribution of all glutamines (Q). Statistical analysis of secondary structure distribution calculated using Chi-squared goodness of fit. H) Score distribution of published cleavage sites (P1=Q), striated by secondary structure. Statistical analysis calculated by one-way ANOVA with Holm-Sidak multiple comparison test. For all statistics shown, * = p ≤ 0.05 , ** p ≤ 0.01, ***p ≤ 0.001, **** = p ≤ 0.00001

Only limited efforts have thus far been taken to identify systematically, and in unbiased fashion, host cleavage targets of 3CLPro from SARS-CoV-2. Given the high conservation of 3CLPro, such analysis would extend as well across coronavirus species. One approach taken recently used N-terminomics to identify neo-N-termini generated by the viral proteases, and identified 14 new cellular substrates ^25^ and more than 100 substrates in a second study ^26^. This approach is limited, however, by the need for sufficient protein abundance and appropriate fragment size and properties to be detected by mass spectrometry ^27, 28^. Only 3 cleavage targets have been identified by more than one study to date (TAB1, ATAD2 and NUP107), reflecting the lack of saturation of these approaches. Moreover, cleaved proteins that are subsequently degraded (a process accelerated by infection) ^29, 30^ also escape detection by N-terminomics, as do proteins not expressed by the cell types used experimentally. *In silico* approaches provide the opportunity to overcome these numerous limitations, and to avoid laborious experimental screens. An initial such approach relied on similarity between cleavage sites in the viral polypeptide across divergent coronavirus species (NetCorona1.0)^31^. However, this method generates scores using assumptions about the viral cleavage site that do not apply to the SARS-CoV-2 consensus sequence. For example, NetCorona1.0 predicts that a sequence containing a proline at the P2’ position can be cleaved, but this substitution has been shown to block cleavage in cleavage assays ^32^. In addition, NetCorona1.0 does not consider cleavage site accessibility conferred by secondary structure, the relative efficiency of cleavage at different sites, or the possibility that there may be host target sites of higher affinity than viral sites.

Here we combine published cleavage efficiency data on the SARS-CoV 3CLPro, which is 96% similar to SARS-CoV-2 3CLpro^33^, with genome-wide secondary structure analyses, to identify and score 99,000+ predicted SARS-CoV/SARS-CoV-2 3CLPro cleavage sites across the human proteome. Through score filtration and secondary structure analysis, we identify over 1000 high likelihood sites. We re-discover nearly all prior SARS-CoV-2 3CLPro experimentally identified sites, and we validate newly identified targets with purified reagents and in cell culture.

Focusing on cardiomyocyte-specific hits, we show 3CLPro leads to cleavage and degradation of the sarcomeric protein obscurin (OBSCN) in human induced pluripotent cell-derived cardiomyocytes (hiPSC-CM), and recapitulates the sarcomeric disorganization observed with SARS-CoV-2 infection of hiPSC-CMs^34–36^. Our study provides a comprehensive atlas for identifying the degradome of 3CL proteases, applicable to SARS-CoV-2 and, in light of the structural conservation of the 3CL protease across coronavirus species ^37^, future coronavirus outbreaks.

### Bioinformatic prediction of SARS-CoV-2 3CLPro targets using experimental data from SARS-CoV 3CLPro

We first sought to identify and score potential cleavage targets of the 3CLpro encoded by SARS-COV-2. Given the 96% sequence similarity between 3CLPro from SARS-CoV-2 as well as the homology in the viral genome cut sites ^17, 33, 37^, we developed an algorithm based on experimental data generated previously from SARS-CoV (2003) 3CLPro^32^. In this previous study, FRET polypeptides spanning the first endogenous cut site between NSP4 and NSP5 (P5-SAVLQSGF-P3’) were generated and modified with every possible single amino acid substitution from P5 to P3’ position relative to the cleavage site. Cleavage efficiency by 3CLpro was then assessed by fluorescence intensity compared to the consensus cleavage sequence. We leveraged this data set to generate a score for every possible cleavage site using a lookup table, multiplying the relative efficiency of each amino acid. This multiplication was then applied with a sliding 8-amino acid windows across the entire human proteome (**Fig 1B**). Substitution at any site that showed no detectable cleavage was interpreted as “0”. Assuming a glutamine (Q) in the P1 position, over 98,697 scored sites (>0) were identified. Expanding our search to include methionine (M) or histidine (H) at P1 uncovered a total of 195,684 sites with a median score of 0.0008 (**Fig 1C**) (**Supplemental Table 1)**. GO analysis of scores in the top 15% (>0.01) showed an enrichment for cell-adhesion, morphogenesis and cytoskeletal genes (**Supplemental Table 2**). We named the algorithm Sarsport1.0.

To evaluate the precision of Sarsport1.0, we calculated scores for the 11 known 3CLPro cut sites in the SARS-CoV viral genome. Scores ranged from 1.31 – 0.04, all within the upper 5^th^ percentile of the score range. These scores were then compared with the published relative K_cat_/K_m_ values for each cleavage site^38^. With the exception of the cut-site between NSP9 and NSP10 (ATVRLQ*AGNAT), our calculated score correlated closely with relative K_cat_/K_m_ (**Fig 1D** left). In contrast, there was essentially no correlation between NetCorona1.0 scores and relative Kcat/Km (**Fig 1D**, right).

To evaluate the sensitivity of Sarsport1.0 to identify SARS-CoV-2 host protein targets, we next calculated scores for the >100 recently published experimentally-identified SARS-CoV-2 3CLPro cleavage targets. Sarsport1.0 identified 104 of 117 sites, including those with non-canonical methionine or histidine at the P1 position (**Fig 1C**). The median score was over 0.1, which is within the top 2.5% of all scores. Receiver operator curve (ROC) analysis (**Fig 1E**) showed Sarsport1.0 to be highly predictive, with an area under the curve of 0.9473, and P < 0.0001. This is likely an underestimate of true ROC, because true positives were likely missed in the experimental approaches. We conclude that Sarsport1.0 is highly predictive of cleavage sites by both SARS-CoV and SARS-CoV-2 3CLpro proteases.

### Refinement of cleavage prediction by secondary structure analysis

The unique high score but low K_cat_/K_m_ of the NSP9/10 cleavage site (**Fig 1D**) suggested that a higher order structure, not captured by scoring based on primary sequence alone, might inhibit cleavage. We therefore estimated the secondary structure of each cut site in the viral genome, using the JPRED4 protein secondary structure prediction server^39^ and a 100aa window spanning the P1 position. The NSP9/10 site in SARS-CoV was the only cleavage site where the P1 position (Q) was predicted to lie in a *β*-sheet (**Supplemental Table 3**). In contrast, the other sites all lay in predicted *α*-helices or disordered regions, structures known to be more accessible to proteases ^40, 41^. These data suggested that higher order structures such as *β*-sheets hinder cleavage by 3CLpro.

To further probe this possibility, we used JPRED4 to evaluate secondary structures around all predicted cleavage sites with a Q at P1 and with a Sarsport1.0 score >0.1 (4416 sites) (**Fig 1F**) (**Supplemental Table 4**). The recent publication of predicted structures for most of the human proteome with AlphaFold^42^ also provides the opportunity to cross validate secondary with higher order structure. The relative frequency of *β*-sheet structures at the P1 position of predicted cleavage sites was significantly less than the overall frequency of *β*-sheets for glutamines in the proteome^43^ (**Fig 1G**), indicating that Sarsport1.0 partly biases away from *β*-sheets. Further comparison to published experimentally-identified cleavage sites revealed in the latter an additional significantly increased propensity for cleavage in regions where P1 (Q) is unstructured and in particular not in a *β*-sheet (**Fig 1F**). Thus, filtering results from Sarsport1.0 for the absence of a *β*-sheet structure at P1 will improve its positive predictive value.

Interestingly, the median Sarsport1.0 score for sites that lie in unstructured regions (0.1024) was significantly lower than for sites that lie in *α*-helices (0.1727) or *β*-sheets (0.27) (**Fig 1G**), suggesting that the presence less permissive secondary structural order imposes a higher evolutionary pressure for an optimal primary sequence cleavage motif.

### Cleavage validation of novel targets

Because of the higher sensitivity of our method, we identified numerous new predicted cleavage sites, in addition to those previously published. Gene Ontology (GO) analysis of proteins with Sarsport1.0 score > 0.01 showed enrichment for many cell-adhesion proteins, including many predicted cleavage sites located within homologous cadherin domains in the cadherin protein superfamily (**Supplemental Table 2)**. Evaluation with AlphaFold predicted these sites to be in unstructured accessible loops within the cadherin domain, thus making them likely to be cleaved if exposed to 3CLPro (**Fig 1G**). We validated these hits *in vitro* by incubation of purified 3CLPro with commercially available recombinant cadherin proteins (CDH6, CDH20), which have identical predicted sites (Score 0.145, Q203 and Q209, respectively, Fig/Table XX). 3CLPro efficiently cleaved both CDH6 and CDH20, yielding the expected fragment sizes based on the predicted cleavage site (**Fig 2A**). We similarly validated novel cleavage sites in thrombin (IIA) and in the intracellular domain of NOTCH1: *in vitro* reactions with purified proteins yielded expected fragment sizes for both (**Fig 2B-D**). The appearance of thrombin IIA cleaved product was inhibited by the 3CLpro inhibitor GC376, demonstrating the requirement of 3CLpro enzymatic activity (**Fig 2B**). Cleavage of NOTCH1 at a predicted site (Q2315, score 0.432), within the intracellular domain of NOTCH1, yielded both predicted fragments (**Fig 2C**). Overexpression of 3CLPro in hiPSC-CMs also yielded NOTCH1 fragments of predicted length, demonstrating cleavage in intact cells (**Fig 2D**). Additional targets chosen for their high score and secondary structure accessibility (SVIL, UACA, NOTCH2) were similarly validated with 3CLPro overexpression in 293T cells, as was the previously published target TAB1 (**Supplemental Fig 1A**). Interestingly, in these cell overexpression experiments, while the levels of full length target proteins were significantly reduced by expression of 3CLpro, the appearance of fragments of predicted size were not observed. We hypothesized that cytosolic fragments generated by 3CLPro may be further degraded by endogenous proteolytic pathways. Supporting this notion, the plasma-membrane bound N-terminal cleavage product of full length NOTCH1 yielded the expected 90kda fragment, while the C-terminal fragment only showed reduction in total protein (**Supplemental Fig 1B**). We conclude that 3CLPro can cleave a wide range of host proteins, and that the generated cytosolic protein fragments are likely often degraded by endogenous pathways.

**Figure 2.**
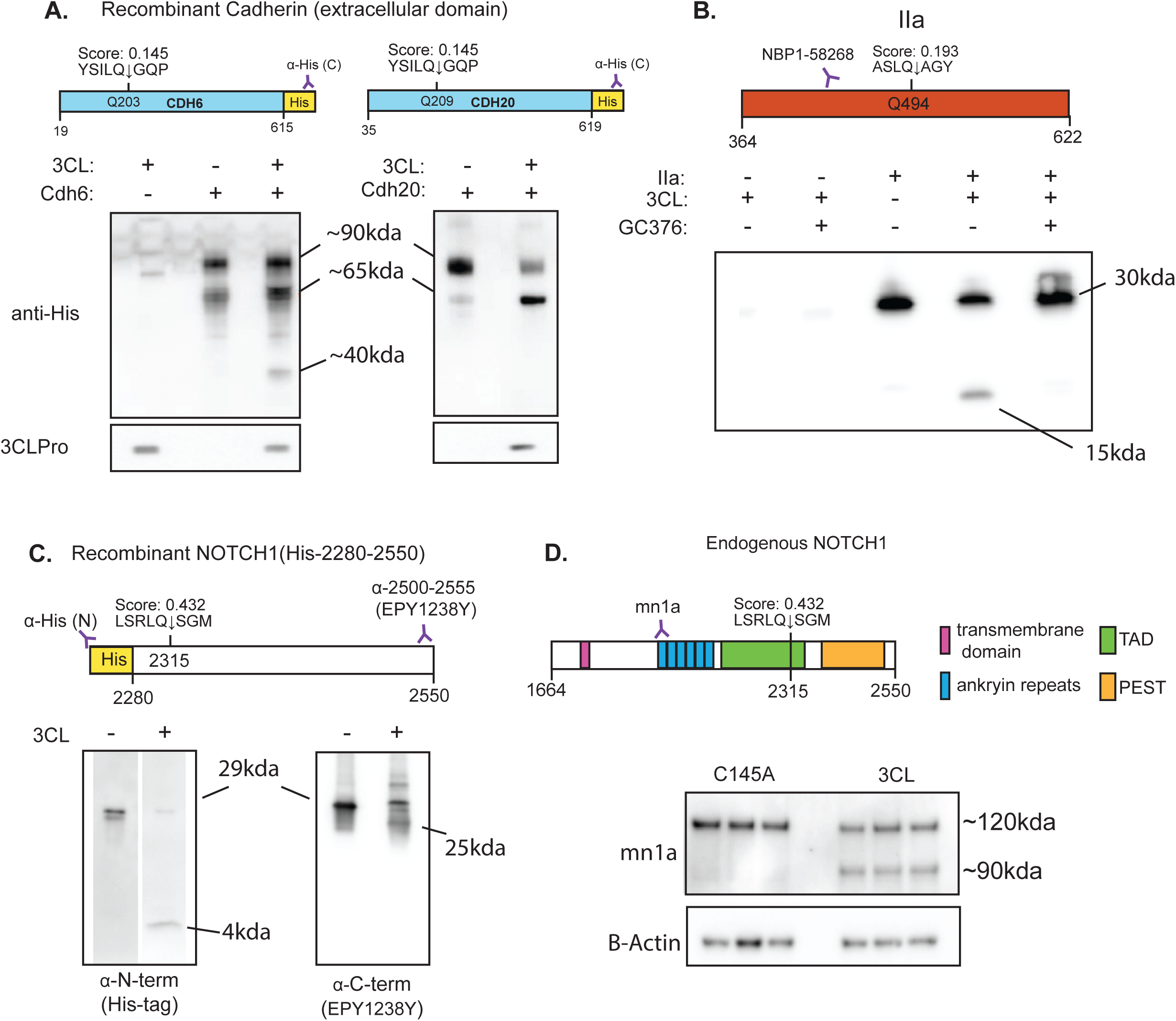
A) Western blot of *in vitro* cleavage of recombinant cadherin with purified 3CLPro. 1uM of purified 3CLPro was incubated with 2ug of recombinant CDH6 or CDH20 (C-terminal His-Tag) in a 50uL reaction for 1hr. Shown are cleavage sites within the recombinant fragment, with amino acid positions displayed for the full length proteins. Western blots showing staining against the C-terminus (His-Tag) of each protein and 3CLPro. Recombinant proteins are a mixture of glycosylated (∼90kDa) and unglycosylated (∼65kda), corresponding to cleavage fragments of ∼62kDa and 40kDa (respectively). B) Western blot of *in vitro* cleavage of purified human alpha thrombin (IIa). Diagram shows amino acid position of unprocessed prothrombin. Position of cleavage site shown with respect to epitope of antibody used for western blot. 1uM of purified 3CLpro was incubated with 2ug alpha thrombin overnight under reducing conditions, with or without the 3CLPro inhibitor GC376 (1uM). C) *In vitro* cleavage of purified recombinant NOTCH1 fragment (aa2280-2550) with a N-terminal His-Tag. Reactions were done with 1uM of purified 3CLPro for 1hr. Diagram shows position of cleavage within the NOTCH1 fragment, with amino acid positions corresponding to the full length protein. Epitope regions showed for antibody with epitope C-terminal to the cleavage site. Full length size is ∼29kDa, with N and C-terminal fragments of 4kDa and 25kDa (respectively). D) Cellular cleavage of NOTCH1. Western blots show lysates of hIPSC cardiomyocytes expressing 3CLPro or catalytically inactive C145A variant for 48h. Cleavage site position within the intracellular fragment of NOTCH1 shown, as well as epitope for antibody used in western blot. For all statistics shown, * = p ≤ 0.05 , ** p ≤ 0.01, ***p ≤ 0.001, **** = p ≤ 0.00001

### Cardiac targets of SARS-CoV-2 3CLPro show multiple cut sites across sarcomeric proteins

Previous work has demonstrated disorganization of sarcomeres after SARS-CoV-2 infection of hiPSC-derived cardiomyocytes ^34–36, 44^. We hypothesized that 3CLPro may be degrading sarcomere proteins directly. Consistent with this notion, overexpression of 3CLPro, but not a catalytically inactive mutant (C145A), in hiPSC-CMs led to pronounced sarcomere breakdown within 48h (**Fig 3A**). At this 48h time point, we also observed numerous cells with a stereotypical intermediate phenotype, in which sarcomeres exhibited increased length, as defined by the distance between alpha-actinin stained Z-discs (**Fig 3B-C**), suggesting that a key structural protein of the sarcomere is being targeted by 3CLpro.

**Figure 3.**
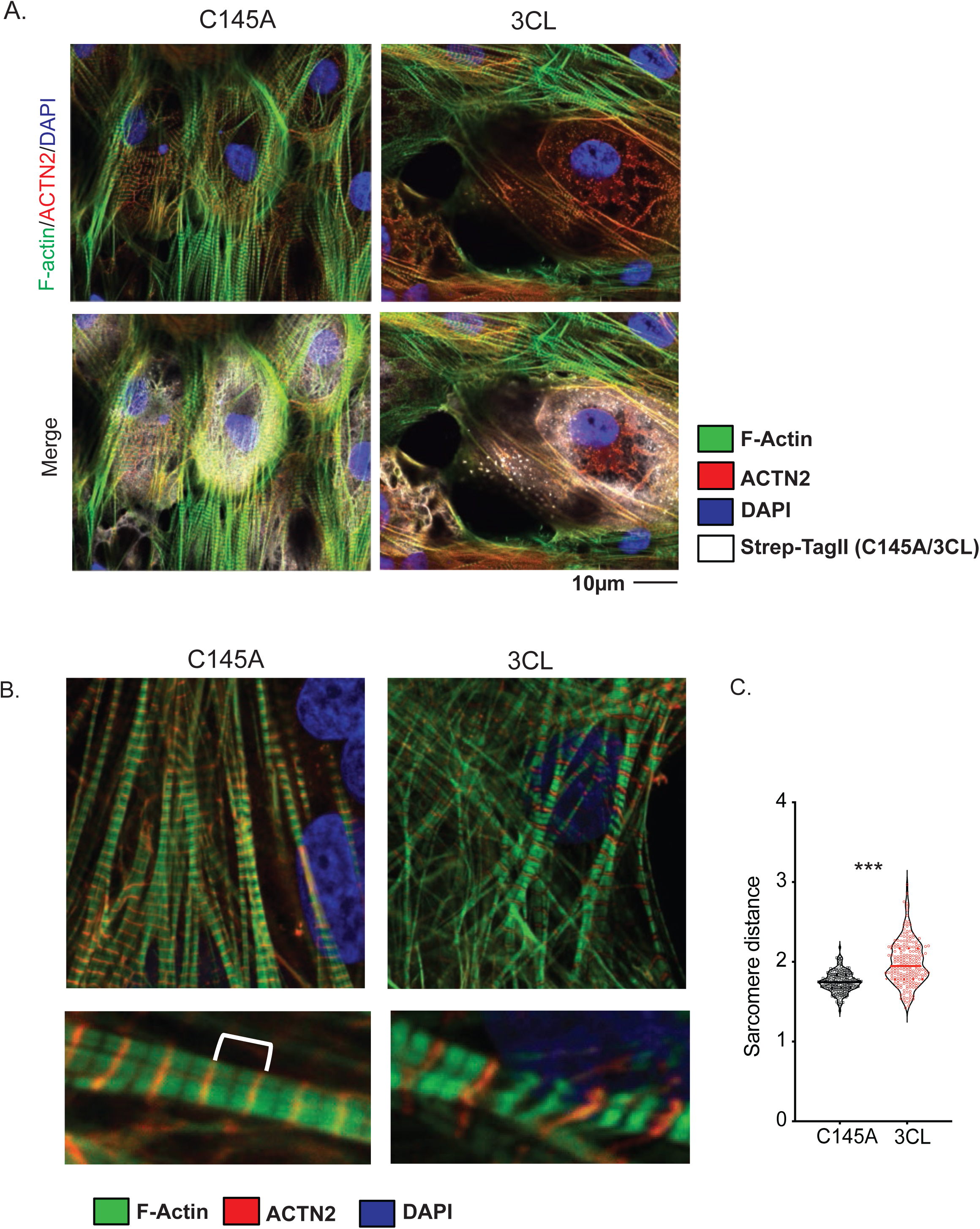
A) Sarcomere breakdown with overexpression of 3CLPro. hiPSC-CMs transduced with adenovirus overexpressing Strep-Tagged 3CLPro or catalytically inactive C145A control for 48h. Staining shows alpha-actinin (ACTN2), F-actin (phalloidin) and Strep-Tag. DAPI counterstain. B) Increased sarcomere length with overexpression of 3CLPro. Sarcomeres are stained with ACTN2 to mark Z-disks. Example image of sarcomeres with increased distance. C) Quantification of sarcomere distance. Sarcomere lengths (Z-disk to Z-disk length, as stained by ACTN2) were quantified for ∼200 sarcomeres (N=2/3 independent experiments? or only 1 exp with 200 sarcomeres?). Statistics shown by Student’s t-test (*** p = XXX). For all statistics shown, * = p ≤ 0.05 , ** p ≤ 0.01, ***p ≤ 0.001, **** = p ≤ 0.00001

We applied our *in silico* primary and secondary analysis to identify putative sarcomere targets of 3CLpro (**Supplemental Table 5**). Within this list, we identified the giant protein Obscurin (OBSCN) as a probable target, with 5 high-likelihood sites along the length of the ∼800kda (>7500 amino acids) protein (**Fig 4A**). Consistent with these data, hiPSC CMs expressing 3CLpro had a marked reduction in Obscurin protein, compared to cells expressing the C145A mutant (**Fig4B-C**). The reduction was apparent using antibodies against multiple epitopes along this large protein. In contrast, levels of alpha-actinin (ACTN2) and myosin heavy chain (MYH6), which our algorithm did not predict to be cleaved by 3CLpro, were not altered (**Fig 4B)**. In addition, immunocytochemistry showed loss of Obscurin in otherwise apparently intact, alpha-actinin-positive sarcomeres in cells expressing 3CLPro, but not cells expressing C145A (**Fig 4C**).

**Figure 4.**
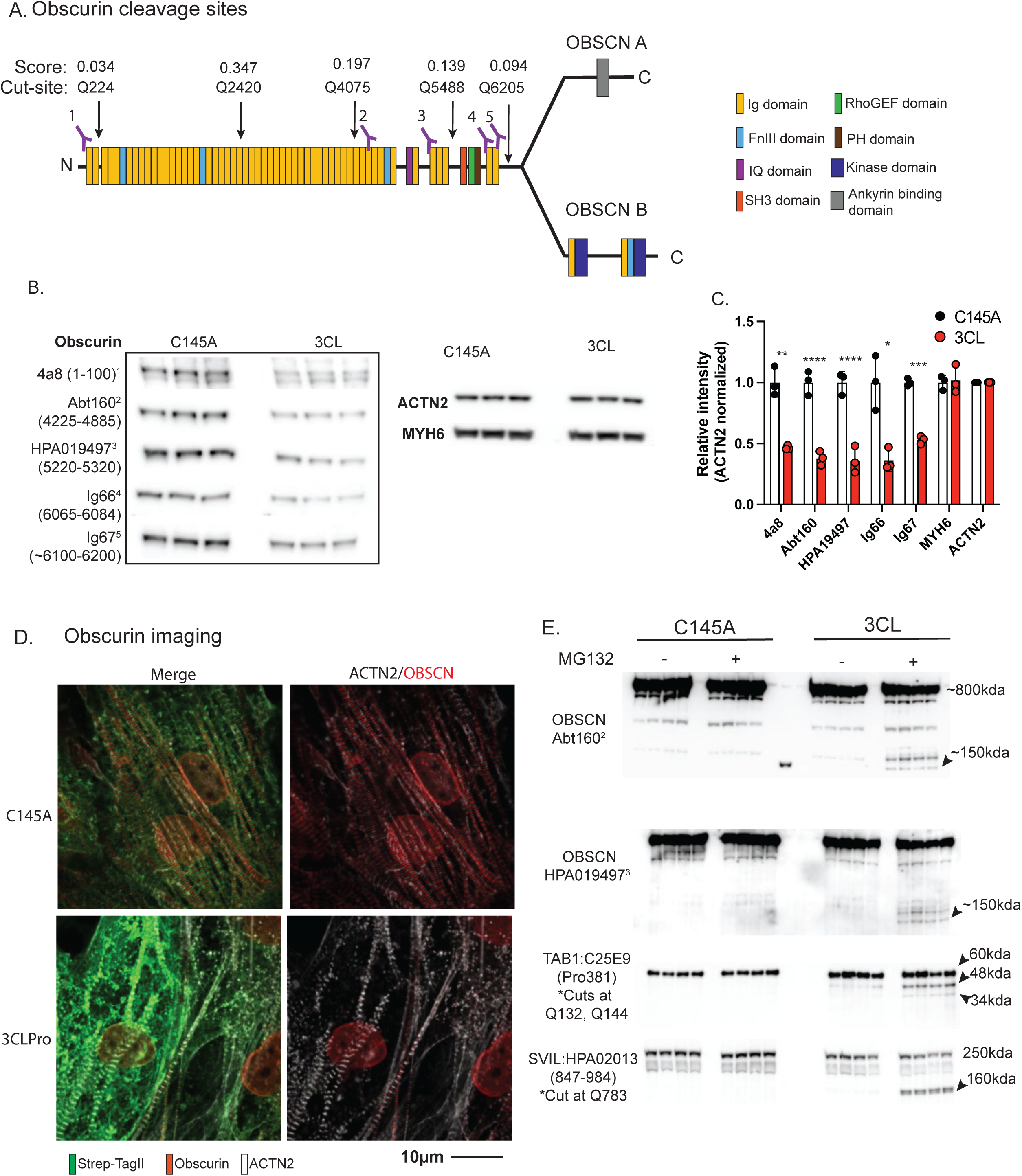
A) Schematic of Obscurin (OBSCN) with overlayed predicted cleavage sites and protein domain. Epitopes regions for antibodies used for this study shown (1-5). B) Obscurin expression after 72h of 3CLPro or catalytically inactive C145A. Western blots shown for all 5 OBSCN antibodies, as well as ACTN2 and MYH6 controls. C) Quantification of blots shown in **B,** normalized by ACTN2 staining on the same membrane. Statistics by Student’s t-test. D) Immunocytochemistry for OBSCN2 and ACTN2 after 48h overexpression of 3CLPro or catalytically inactive C145A. E) Expression of cleavage targets following expression of 3CLpro or catalytically active C145A. Following 24h of overexpression, MG132 (1uM) or vehicle was added for 24h for a total of 48h overexpression with or without MG132. Fragments are highlighted for OBSCN, TAB1, and SVIL. For all statistics shown, * = p ≤ 0.05 , ** p ≤ 0.01, ***p ≤ 0.001, **** = p ≤ 0.00001

As with a number of targets described above (**Supplemental Fig 1**) we did not detect any Obscurin fragments, despite using antibodies that recognize multiple epitopes along the length of the protein. To test whether the absence of fragments might be due to endogenous proteosome activity, we treated cardiomyocytes expressing 3CLPro, versus C145A, with the proteosome inhibitor MG132, and collected cellular protein 24h later. Blotting with the two antibodies that recognize the region between cut sites 3 and 4 (Q4075 and Q5488, respectively) yielded the expected fragment size (∼155kda) (**Fig 4D**), validating the predicted sites as a 3CLpro targets, and demonstrating that the ensuing fragment is targeted for degradation by the proteosome. Identification of other fragments within Obscurin was technically unfeasible due to either lack of antibodies against the specific region, or to overlap with non-specific bands on blots. However, we also observed the appearance, after proteasome inhibition, of fragments of predicted size in Supervillin (SVIL), another giant sarcomeric protein predicted to be targeted by our algorithm, as well as TAB1, which was previously shown to be targeted by 3CLpro but for which no fragments had been detected^24^

### Obscurin degradation in SARS-CoV-2 infection

Finally, we tested if those results were recapitulated in a model of hPSC-CMs infected with live SARS-CoV-2. For these experiments, we used two hPSC-CM lines, WTC-11c (hiPSC-CMs) and H7 (hESC-CMs, human embryonic stem cell-derived cardiomyocytes), previously used to study the effects of SARS-CoV-2 on human cardiomyocytes^35^. Within 48h of infection, coincident with the robust appearance of viral nucleocapsid, total Obscurin protein gradually reduced by 40-60% (**Fig 5A-B**). In contrast, protein abundance of ACTN2, MYH6, and MYH7 was unaffected by infection (**Fig 5B**), mirroring the effects seen with 3CLpro alone (**Fig 4B**). Similarly, immunocytochemistry of infected cells showed reduction of Obscurin staining in cells expressing viral nucleocapsid, despite seemingly intact sarcomeres (TNNT2 staining), again mirroring the effects seen with 3CLpro alone **(Fig 4D**). Thus the loss of Obscurin caused by 3CLpro-mediated cleavage might explain the loss of sarcomere integrity during SARS-CoV-2 infection in human cardiomyocytes.

**Figure 5.**
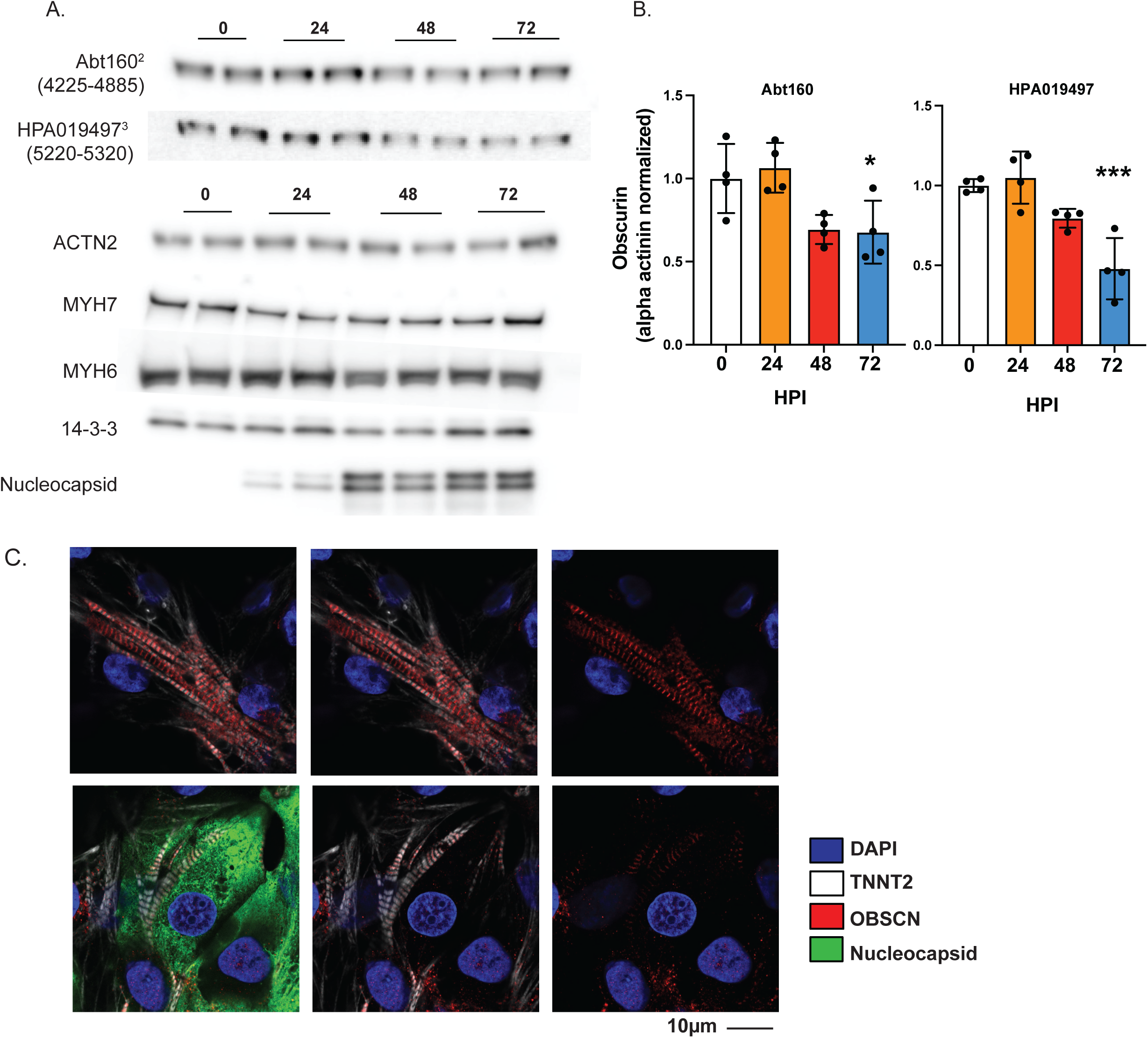
Obscurin degradation in SAR-COV2 infection. A) Western blots of WTC-11c hPSC-CMs infected with SARS-CoV-2 at 5 MOI (Multiplicity of infection) after 24, 48 and 72h. B) Quantification of OSCN staining normalized for ACTN2 (per blot). Shown are n=2 for each time point for 2 different hPSC-CMs cell lines (WTC-11c and H7) for a total of n=4. Statistics calculated by one-way ANOVA with Tukey’s post-hoc test. C) Immunocytochemistry for OBSCN in WTC-11c hPSC-CMs at 48 HPI with 5 MOI SARS-CoV-2. TNNT2 used as a counterstain for sarcomeres, and nucleocapsid staining performed to identify infected cells. For all statistics shown, * = p ≤ 0.05 , ** p ≤ 0.01, ***p ≤ 0.001, **** = p ≤ 0.00001

## Discussion

We leveraged here experimental data and genome-wide secondary structure analyses to develop a reliable computational algorithm, Sarsport1.0, that predicts endogenous non-viral cleavage targets by the 3CLpro SARS-CoV-2 protease across the human proteome. We validated the precision of the algorithm by confirming novel predicted cleavage sites, using both biochemical and cell culture approaches. The algorithm is specific: all 8 predicted host proteins that we chose to test experimentally confirmed cleavage by 3CLpro. The algorithm is also precise, accurately correlating scores with the known K_m_/K_cat_ values for the cut sites in the single viral polypeptide. The single exception to this correlation, the NSP9/10 site, is also the only site lying within a predicted beta-sheet, which likely hinders protease access. The particularly high score of the NSP9/10 site (score = 1.31) may have evolved to overcome this more inaccessible higher order structure.

Finally, the algorithm is highly sensitive, accurately predicting high cut scores in nearly all previously experimentally identified 3CLpro sites ^23–26^. In addition, thousands of novel sites are predicted to be cleaved by 3CLpro. Numerous reasons likely explain why the computational algorithm is more sensitive than prior experimental approaches: (1) experimental paradigms are limited to detecting the proteins expressed in the chosen experimental cells; (2) proteomic approaches rely on the ability to detect new protein fragments by mass spectrometry, a relatively insensitive method; (3) and proteomic approaches also require the physical presence of cleaved products, but as shown here, these fragments are often quickly degraded after cleavage. Our computational algorithm overcomes these limitations and provides a comprehensive atlas of the putative SARS-CoV-2 3CLpro degradome.

The SARS-CoV-2 3CLpro is the target of ongoing therapeutic efforts to treat COVID-19. Recent interim analysis of the phase 2/3 EPIC-HR (Evaluation of Protease Inhibition for COVID-19 in High-Risk Patients)^22^, which tested the combination therapy of PF-07321332, a 3CLpro inhibitor, with ritonavir, a CYP3A4 inhibitor that prevents the metabolism of protease inhibitors, reported a nearly 90% reduction in hospitalization or death compared to placebo in non-hospitalized high-risk adults with COVID-19 (P<0.0001). Our data suggest that these remarkable benefits of 3CLpro inhibition may accrue from effects beyond suppression of viral replication. For example, infected cells that do not sustain replication may nevertheless experience significant cellular damage from 3CLpro activity on the host proteome, and PF-07321332 and other 3CLpro inhibitors would be predicted to prevent this cellular damage. Similarly, expression of 3CLpro is one of the earliest events in the viral life cycle, and may thus cause early cellular damage, potentially suppressing cellular defenses against the ensuing viral replication. Lingering effects of 3CLpro may also contribute to persistent symptoms, as observed with the long-COVID syndrome. In sum, the remarkable benefits of 3CLpro inhibition in COVID-19 patients underscores the need to further understand the impact of 3CLpro on the host proteome, which will be substantially aided by our predictive algorithm.

To validate our algorithm, we investigated the effects of 3CLPro in two different models (OE and live SARS-CoV-2) on hiPSC-CMs, previously shown to be susceptible to SARS-CoV-2 infection with profound effects on sarcomeric organization ^34–36^. We identified the giant sarcomeric protein Obscurin as a target of 3CLpro, and showed (1) that both ectopic expression of 3CLpro and infection by SARS-CoV-2 cleave and degrade Obscurin, while leaving other sarcomeric proteins intact; and (2) that ectopic expression of 3CLpro, but not a mutant without enzymatic activity, causes sarcomeric disorganization in a stereotypical fashion similar to that observed with SARS-COV-2 infection, and consistent with the pattern predicted by degradation of Obscurin, an important structural component of the Z-disk. Thus, we propose that sarcomeric disorganization during SARS-CoV-2 infection is likely caused in large part by direct proteolysis of Obscurin by 3CLpro.

Cardiac complications of COVID19 have been well documented, and the presence of cardiac damage, as reflected in plasma troponin levels, is highly predicted of morbidity and mortality after SARS-COV-2 infection ^45–47^. It is still unclear if the pathology observed in the heart is due to a direct cytotoxic effect of the virus or secondary to the systemic inflammation. Some evidence of direct infection by SARS-CoV-2 in the human heart has been reported, suggesting that a direct effect is possible^48, 49^, although other post-mortem studies have not detected infected cardiomyocytes in COVID-19 patients^44–46^. Thus, the implications of our findings in iPSCMs on human cardiac disease should be interpreted cautiously. We used these studies primarily as molecular and structural validation of our predictive algorithm. The use of hPSC-CMs also offers a powerful tool for pharmacological and drug screening, in addition to disease modeling.

In summary, we provide a validated, computationally derived, comprehensive atlas of the putative 3CLpro degradome, overcoming limitations in sensitivity inherent to experimental approaches. Our findings provide a powerful tool to aid investigations into the virus-host interactions mediated by 3CLpro, the target of highly efficacious therapy against COVID-19. In light of the structural conservation of the 3CL protease across coronavirus species, such investigations will likely also apply to future coronavirus outbreaks.

## Acknowledgements

ZA was supported by NIH/NHLBI (HL152446). NY was supported by NIH training grant 5T32AR053461. Q.M was supported by NIAMS T32 training grant (AR 53461-12). SM was supported by post-doctoral fellowship from the Institute for Stem Cells and Regenerative Medicine (University of Washington). This work was also supported by R01 HL128362 and R01 HL146868 to C.E.M.; the Robert B. McMillen Foundation and the State of Washington philanthropical support to the UW Institute for Stem Cell and Regenerative Medicine. We would also like to acknowledge the staff of the University of Washington BSL-3 facility, as well as our Environmental Health and Safety staff, who provided assistance during experiments and ensured our safety.

---

**Supplementary Figure 1.**
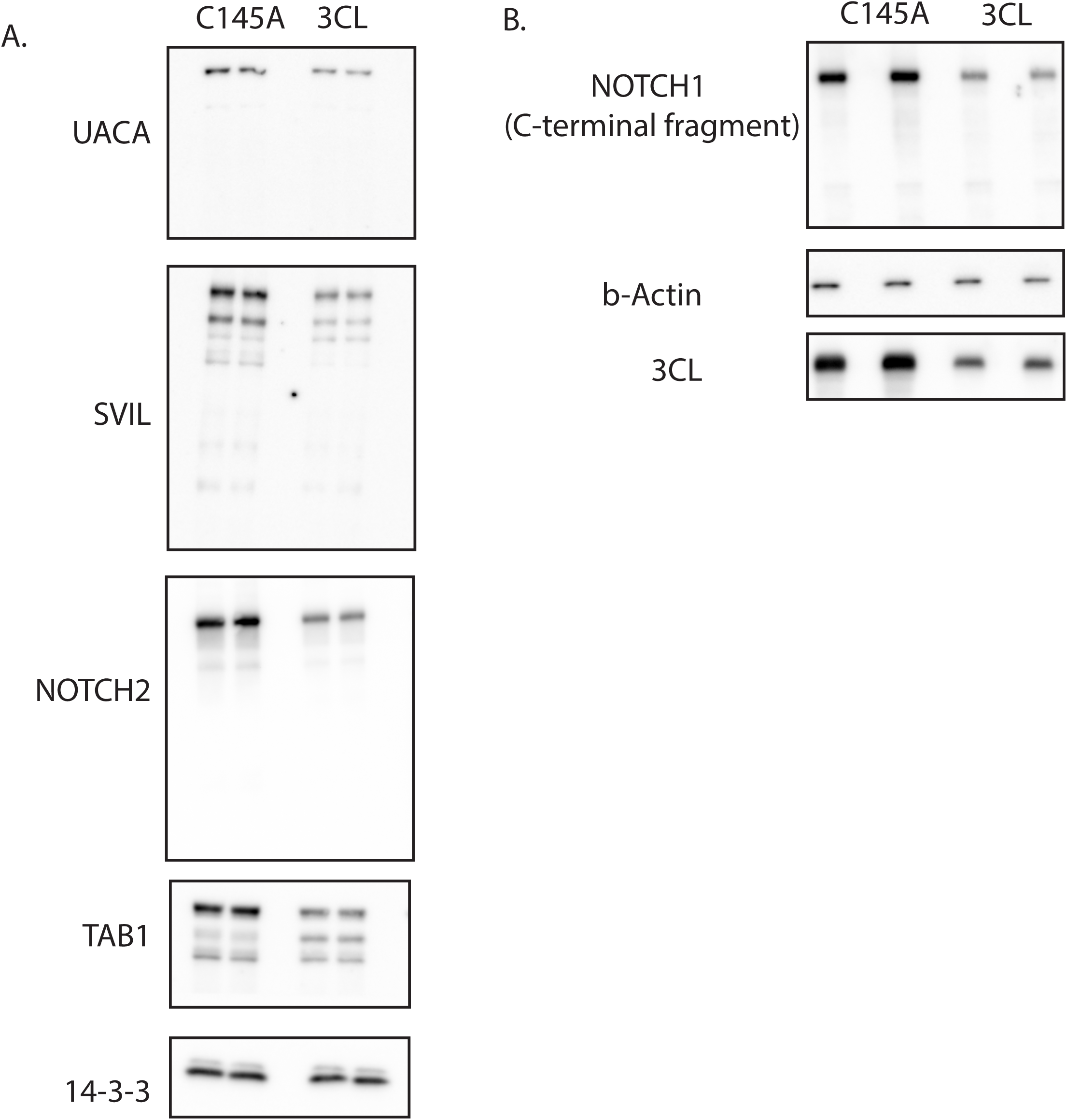
A) Western blots of candidate cleavage targets. 48h overexpression of 3CLPro or C145A in 293T cells. B) Western blot for the C-terminal cleavage fragment of NOTCH1 following 48h overexpression in 293T cells.

## Methods

### Bioinformatic prediction of 3CLpro cleavage sites

In brief, P1 positions (Q, M or H) were first identified across the human proteome. 8 amino acid peptides were generated centered at this position, corresponding to P5 – P3’ positions in the 3CLPro consensus sequence. Scores were generated by multiplying the relative efficiency values published by Chuck, et al for the SARS-COV (2003) 3CLPro. All sites with a score > 0 (i.e., those that did not have an “ND” in any of the positions from P5-P3’) were captured. The full code for this program will be shared upon publication.

### In vitro cleavage assays

In vitro cleavage assays were performed with purified 3CLPro protein and assay buffer from BPS Bioscience. Protease was added at 1uM concentration, and recombinant protein targets at an approximate ratio of 2ug target/1ug 3CLPro. Recombinant proteins were purchased commercially: NOTCH1 (Origene, Cat# TP762041), CDH6 (ACROBiosystem, Cat# CA6-H5229), CDH20 (R&D, Cat# 5604-CA-050). Purified human alpha-thrombin was purchased from Haematologic Technologies (Cat# 50-883-435).

### Cell Culture

For 3CLPro overexpression assays, human induced pluripotent (hIPSC) ventricular cardiomyocytes were purchased from NCardia. Cells were plated at ∼150,000 cells/24-well on fibronectin coated (Sigma, F1131), glass bottomed nano-patterned plates or coverslips (CuriBio, ANFS-0024). Following 4 days of maturation, cells were transduced with adenovirus (Vector Biolabs) to induce expression of 3CLPro or catalytically inactive C145A, and lysates collected at 48 or 72h in TU buffer.

### Human induced-pluripotent stem cells culture and differentiation

Human induced-pluripotent stem cells (WTC11c hiPSCs, gifted by Dr. Bruce Conklin, Gladstone Institutes, San Francisco) were maintained in complete mTeSR Plus (Stem Cell Technologies) and cultured on Matrigel-coated dished at 0.17mg/mL (Corning). WTC hiPSCc were passaged as small clumps for maintenance or single cell-like suspension for cardiac differentiation using Versene (Gibco) and mTeSR Plus supplemented with 10 μM Y-27632 (Tocris). Cardiac differentiation was perfomed as previsoudly described (PMID: 33657418). Briefly, WTC hiPSCs were seeded at 1,000 cells/cm^2^ using mTeSR1 Plus and 10 μM Y-27632 on Matrigel-coated dishes. After 24 h, media was replaced with mTeSR Plus supplemented with 1 µM Chiron 99021 (Cayman) to prime the cells for differentiation. Mesoderm induction (Day 0) was performed with 3 µM Chiron 99021 in RPMI-1640 media (ThermoFisher) supplemented with 500 μg/mL BSA (Sigma-Aldrich) and 213 μg/mL ascorbic acid (Sigma-Aldrich), named RBA media. After 48 h (Day 2), cells were treated with RBA media supplemented with 2 µM WNT-C59 (Selleckchem). On day 4, media was change with RBA only and cells were incubated for an additional 48 h.

From day 6 until day 13, hiPSC-derived cardiomyocytes (hiPSC-CMs) were maintained in RPMI-1640 supplemented with B-27 supplement (ThermoFisher). Heat-shock was performed at 42C for 30 min and on day 14, hiPSC-CMs were dissociated using 0.5% Trypsin (Gibco) and cryopreserved in Cryostore CS10 (Sigma).

### HiPSC-CMs infection

All experiments using live virus were performed in the Biosafety Level 3 (BSL-3) facility at the University of Washington in compliance with the BSL-3 laboratory safety protocols (CDC BMBL 5th ed.) and the recent CDC guidelines for handling SARS-CoV-2. Before removing samples from BSL-3 containment, samples were inactivated by Thiourea buffer or 4% paraformaldehyde, and the absence of viable SARS-CoV-2 was confirmed for each sample by plaque assays described in the next section.

Frozen hiPSC-CMs were thawed in RPMI 1640 supplemented with B27 supplements, 10 µM Y-27632 and 5% FBS. After 24 h, media was replaced with RPMI 1640 supplemented with B27 supplements only. A total of 200.000 hiPSC-CMs were seeded three days after thawing in Matrigel-coated 24-wells plate using RPMI 1640 supplemented with B27 supplements, 10 µM Y-27632 and 5% FBS. Media was changed after 24 h and infection was performed as previously described (PMID: 33657418). Briefly, hiPSC-CMs were quickly washed with DPBS and incubated with SARS-CoV-2 at 5 multiplicity of infection (MOI) diluted in DMEM only (Gibco) for 1 h at 37 C. Mock-control hiPSC-CMs were treated with DMEM only. Media was then replaced with RPMI 1640 supplemented with B27 supplements and samples were collected at 48 h after infection. For MG132 treated samples, after viral absorption, media was replaced with RPMI 1640 supplemented with B27 supplements and 1 µM of MG132 (DMSO was used for control).

### SARS-CoV-2 preparation and titer

SARS-Related Coronavirus 2, Isolate USA-WA1/2020 (SARS-CoV-2) was obtained from BEI Resources (NR-52281). Virus propagation and titer was performed in VERO cells (USAMRIID) as described in PMID: 33657418. Briefly, VERO cells were maintained in DMEM supplemented with 10% FBS, 100 U/mL penicillin, and 100 U/mL streptomycin and incubated with either 0.1 MOI (virus propagation) or serial dilution of conditioned media (titer) for 1 h at 37C in DMEM only media for viral absorption. For viral propagation, conditioned media was harvested and aliquots were store in -80C. For titer, 10-fold serial dilutions of conditioned media (either from VERO cells or hiPSC-CMs) were incubated on VERO cells for 1 h at 37C. A 1:1 mixture of cellulose suspension (Sigma) and DMEM containing 4% heat-inactivated FBS, L-glutamine, 1X antibiotic-antimycotic (Gibco), and 220 mg/mL sodium pyruvate was layered on top of the cells and incubated at 37C for 48 h. Cellulose layer was then removed and cells were stained with 10% paraformaldehyde and stained with 0.5% crystal violet solution in 20% ethanol. Plaques were counted, and the virus titer in the original sample was assessed as plaque-formation unit per mL (PFU/mL).

### Lysate preparation and Western blotting

For western blotting of 293T cell lysates, traditional RIPA buffer was used to lyse cells. Lysates were normalized for cell concentration by BCA assay, and denatured by addition of Laemmli buffer (BioRad) at 95C for 5 minutes. Samples were then run on tris-glycine gels, followed by transfer onto PVDF membranes. Samples were blocked for 45 minutes with 5% milk in TBS + 0.1% Tween-20, and stained overnight at 4C in Pierce protein-free blocking buffer (Thermo-Scientific).

For cardiomyocyte lysates, cells at equivalent densities were first lysed in thiourea denaturing buffer (TU buffer: 8M urea, 2M thiourea, 50mM Tris-HCl pH 7.5, 3% SDS, 75mM DTT). Following incubation in TU buffer for 5 minutes, an equivalent volume of 50% glycerol was added to lysates for a final concentration of 4M urea, 1M thiourea, 25mM Tris HCl pH 7.5, 1.5% SDS, 25% glycerol and 37.5mM DTT and stored at -80C until western blotting. Samples were not heated so as to avoid urea decomposition, and subsequent carbamylation of proteins from cyanate ions. Phenol red powder was added directly to lysates for visualization These denaturing, highly reducing conditions were crucial for solubilization of sarcomeric proteins. Lysates were run directly on tris-glycine or tris-acetate gels, transferred to PVDF membrane, and stained as described above.

### Imaging

Imaging was performed on a Zeiss LSM 710 confocal microscope at 63X. For sarcomere quantification, lengths of 400 sarcomeres were quantified for n=3 biological replicates. Images assignments were blinded during quantification.

### Antibodies

The following antibodies were used for western blotting. Dilutions were 1:1000 unless specified otherwise.

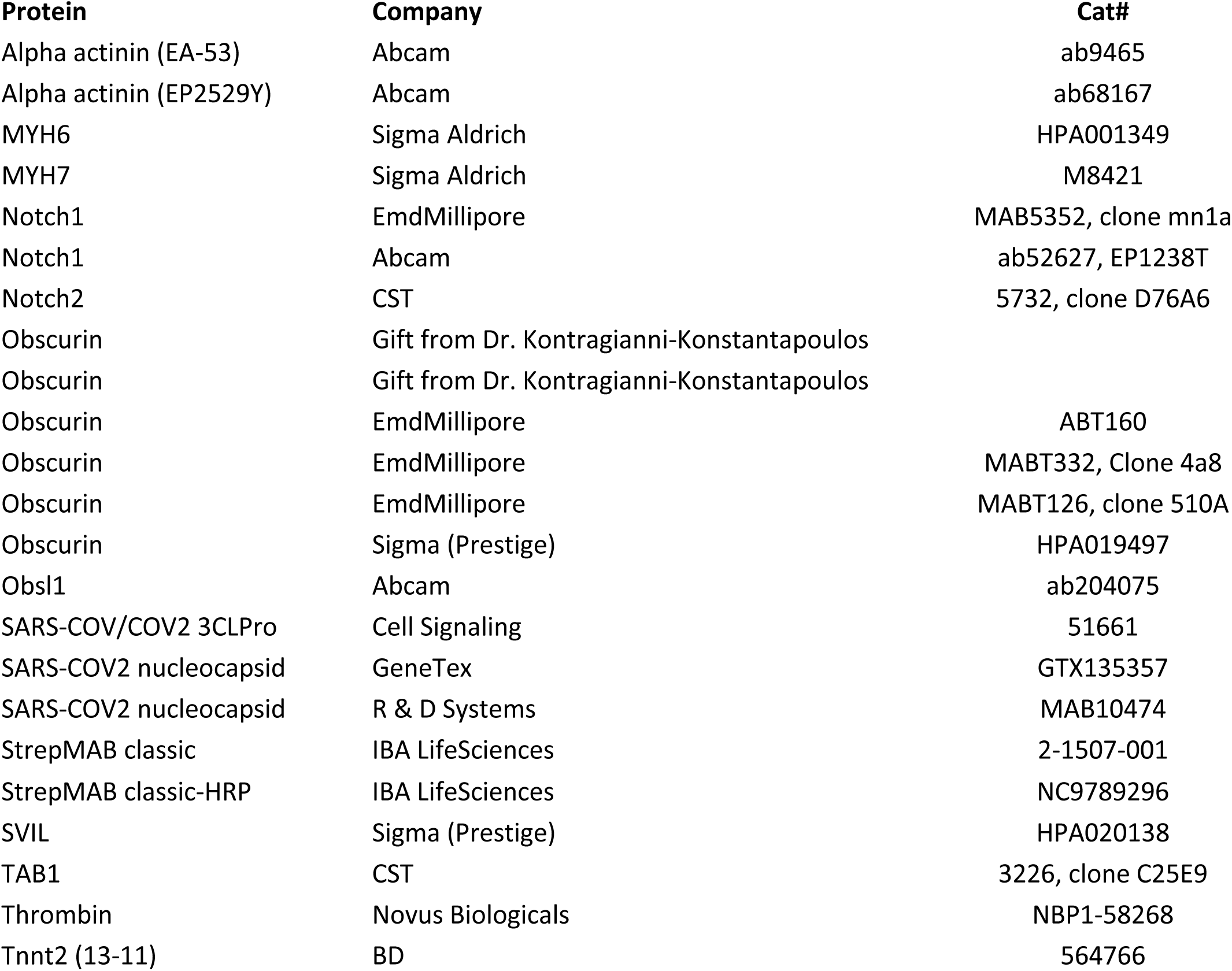
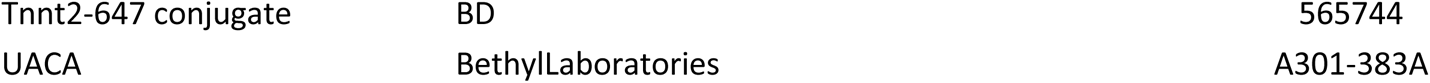

### Statistics

Statistics were calculated with Prism9.

